# Dynamic interaction between stress hormones and neutrophils promotes neutrophil extracellular trap formation with behavioral consequences

**DOI:** 10.1101/2025.11.12.688017

**Authors:** Kannan Thangamani, Helder Prece, Julia Carter, Orion Douglas, Eric Brengel, Craig Ferris, Emeka Okeke

## Abstract

Recent studies have highlighted the crosstalk between neuroendocrine responses and the immune system but the mechanisms underlying this cooperation are still not well understood. The stress response is associated with peripheral inflammation suggesting that stress hormones including glucocorticoids and catecholamines could modulate the function of innate immune cells like neutrophils. Likewise, inflammatory mediators produced by immune cells are known to contribute to psychiatric diseases like major depressive disorder. Here we investigated the dynamic relationship between stress hormones and neutrophils and their contribution to mood disorders. We found that chronic restraint stress leads to plasma elevation of neutrophil extracellular traps (NETs) and increased NET formation in mice. Interestingly, the stress hormones, cortisol and epinephrine induce NET formation in human neutrophils ex vivo. Activation of neutrophils to form NETs leads to their increased expression of adrenergic and glucocorticoid receptors and neutrophil production of both cortisol and epinephrine indicating an autocrine/paracrine mechanism for the regulation of neutrophil inflammatory response by stress hormones. Strikingly, administration of NET components to mice induces depressive-like behavior. Moreover, activation of the glucocorticoid receptor in human volunteers leads to increase in gene expression of NET proteins. Furthermore, patients with major depressive disorder show gene upregulation of NET proteins. Our data highlights the bi-directional relationship between neuroendocrine processes and neutrophils that contribute to stress-induced increase in inflammation and the role of neutrophil inflammatory responses in propagation of behavioral changes following stress.

## Introduction

Chronic stress in daily life is a major risk factor for the development of psychiatric diseases like major depressive disorder (MDD) (1–3). Depression is the single most common mental disorder in the United States (4). The prevalence of depression in US adults is up to 20% with significant economic burden and major contribution to disability, morbidity and mortality (5). Stress activates the hypothalamic-pituitary-adrenal (HPA) axis leading to the release of the stress hormone cortisol (6). Increased levels of cortisol has been reported in depressed patients (7) and prolonged cortisol exposure leads to structural changes in the brain and subsequent behavioral disorders (8). Stress also leads to the activation of the sympathetic nervous system (SNS) by the hypothalamus leading to the production of epinephrine (9) and increased levels of epinephrine is also associated with psychopathology (10, 11).

Recent studies implicate neuro-immune interactions in the pathogenesis of stress-induced psychiatric disorders (12–14). This bidirectional relationship between stress and the immune response is much less understood but emerging evidence indicates that it is a critical determinant of the psychopathology of stress on the individual. Hence, there is critical need to delineate the relationship between stress response and immune response.

The effect of cortisol on the immune response has been shown to be context-dependent with initial studies suggesting a pro-inflammatory effect of cortisol in acute stress (15, 16) and an immune suppressive effect of cortisol during chronic stress (17, 18). However, recent findings indicate that long-term cortisol exposure is also associated with a proinflammatory phenotype (19). Notably chronic stress results in glucocorticoid receptor resistance which leads to failure to down-regulate inflammatory response (20). Prolonged stress leads to systemic immune activation and inflammatory cytokine production (21–23) and cortisol has been shown to enhance the production of pro-inflammatory cytokines by activated monocytes (24).

Similarly, epinephrine has been shown to exert complex immunomodulatory functions. Indeed, several immune cells express β2-adrenergic receptors (25–27) and epinephrine has been shown to influence immune cell distribution, causing a rapid mobilization of monocytes and neutrophils into the bloodstream (28). Also, injection of beta-adrenoceptor agonist has been shown to induce inflammatory cytokine production in rats (29) and epinephrine has been shown to enhance the immune responses of lipopolysaccharide (LPS)-stimulated macrophages (30).

Neutrophils are the most abundant leukocytes in circulation accounting for about 60% of the leukocyte population (31). They are the first responders to pathogenic insult and are critical for immune defense as shown by the susceptibility of neutropenic patients to infections (32).

Mechanisms of host protection by neutrophils include production of cytokines and chemokines, production of reactive oxygen species (ROS) and phagocytosis (33). Neutrophils express glucocorticoid and adrenergic receptors and both cortisol and epinephrine have been shown to modulate neutrophil functions including neutrophil trafficking, recruitment and infiltration during inflammation (34–38). Indeed, neutrophils have been shown to produce epinephrine which contributes to inflammation pathology (39).

As part of their inflammatory response, neutrophils extrude their cellular contents in a framework of DNA in a process called neutrophil extracellular trap (NET) formation (40). Although NETs were first shown to protect the host from infection by containment and killing of microbes (40), subsequent studies unraveled the pathological role of NETs and NETs have been implicated in the pathogenesis of several autoimmune and infectious diseases (33, 41–43). Recent report that NETs increase in the plasma of patients with schizophrenia and that this increase is associated with early life adversity suggests that psychological stress induces NETs formation and that stress-induced NETs contribute to the development of mental health disorders (44).

Here we investigated the bi-directional relationship between stress and NETs formation and the possible contribution of NETs to the development of mental health disorders. We found that chronic restraint stress in mice leads to increased levels of NETs in the circulation and promotes NETs formation in murine neutrophils. Both cortisol and epinephrine induce NETs formation in cultured human neutrophils. Neutrophils activated to form NETs release cortisol and epinephrine and have increased expression of adrenergic and glucocorticoid receptors indicating an autocrine/paracrine mechanism for the modulation of neutrophil inflammatory response by stress hormones. Importantly, we show that injection of NETs leads to depressive-like behavior in mice. Furthermore, activation of the glucocorticoid receptor in humans upregulates the gene expression of NET proteins. Our results highlight the bi-directional mechanism by which stress and NETs formation contribute to the development of psychiatric disorders.

## Materials and Methods

### Animal Work

All animal procedures were reviewed and approved by the Institutional Animal Care and Use Committee (IACUC) at Northeastern University and conducted in accordance with the National Institutes of Health (NIH) *Guide for the Care and Use of Laboratory Animals*. Male and female C57BL/6J mice (strain #000664, Jackson Laboratory, USA) aged 8 weeks were used. Animals were housed in a temperature- and humidity-controlled barrier facility with food and water available *ad libitum*. A reverse 12-h light/dark cycle (lights on 9:00 p.m.–9:00 a.m.) was used to align experiments with the active phase of the circadian cycle. Mice were acclimated for at least one week before experimentation.

### Chronic restraint stress model

Mice were placed in custom restrainers constructed from 50 mL Falcon tubes perforated with air holes for ventilation. Restrainers restricted lateral movement but allowed limited forward–backward motion. Mice were restrained for 3 h/day during their active phase for up to 21 days. Control animals remained undisturbed in home cages.

For corticosterone measurement, blood was collected by tail snip between 7am and 9am on day 0, 14 and 21 using lithium heparin–coated capillaries (240902, Greiner Bio-One, Monroe, NC, USA). Plasma (diluted 1:10) was analyzed with a corticosterone ELISA kit (RE52211, IBL, Hamburg, Germany). For immunity studies, mice were sacrificed and spleen, femur and tibia were collected.

NETs were injected via intraperitoneal route. A cocktail of major NET proteins was prepared by combining 500 μg DNA (J64400.MC, Thermo Fisher, Waltham, MA, USA), 500 μg elastase (0000428612, Sigma-Aldrich, St. Louis, MO, USA), and 2 mg histone (0000423842, Sigma - Aldrich, St. Louis, MO, USA) in PBS, followed by 10-s pulsed sonication.

### Neutrophil isolation

Blood was collected from the peripheral vein of healthy volunteer donors after written informed consent under an IRB-approved protocol at Northeastern University. Samples were collected into heparinized tubes (367874, BD Vacutainer, Franklin Lakes, NJ, USA) and neutrophils were obtained by first spinning the blood on Ficoll-Paque (10355936, Cytiva, Marlborough MA, USA) then subjecting the red blood cell (RBC) layer to 1.5% dextran sedimentation, followed by hypotonic lysis as we previously described (41). Isolated neutrophils were washed with PBS before use. Neutrophils from mouse blood were obtained using a neutrophil isolation kit (19762, Stemcell Technologies, Vancouver, BC, Canada).

### NET induction, quantification and visualization

NETs were induced, quantified and visualized as we previously described (41). Briefly, isolated neutrophils were resuspended in RPMI 1640 medium supplemented with 3% fetal bovine serum (NET medium). The cells were seeded at 1 × 10 cells/well in either 96-well plates (165305, Thermo Fisher, Waltham, MA, USA) or chamber slides (06032440, Lab-Tek, Thermo Fisher, Waltham, MA, USA) and activated for the said duration with phorbol 12-myristate 13-acetate (PMA; P8139, Sigma-Aldrich, St. Louis, MO, USA), cortisol (H0888-5G, Sigma-Aldrich, St. Louis, MO, USA) or epinephrine (204400010, Thermo Fisher, Waltham, MA, USA) at 37 °C.

NETs released in culture were quantified by measuring the fluorescence intensity of extracellular DNA using 1 μM Sytox Green (S7020, Invitrogen, Waltham, MA, USA) for 5 min. For microscopy studies, cultured cells were fixed, stained with sytox green or neutrophil elastase antibody (sc55549, Santa Cruz Biotechnology, Dallas, TX, USA) and imaged using a Zeiss LSM 710 confocal or Zeiss widefield microscope (Carl Zeiss Microscopy, Jena, Germany). For each condition, 16–25 random fields were captured. Images were batch-processed in ImageJ: converted to 8-bit, thresholded (Triangle algorithm), and analyzed for particle size (>68 pixels²) and circularity (0.00–0.80). Quantified overlays were exported for analysis. Elastase activity was measured using (Z-Ala-Ala-Ala-Ala)_D_ Rh110 substrate (11675, Cayman Chemical, Ann Arbor, MI, USA). Standards (10 mU/mL serial dilutions) and plasma samples (diluted 1:10) were incubated with substrate at 37 °C for 30 min. Fluorescence was read (Ex 485 nm/Em 525 nm).

### Stress hormone measurements

Culture supernatants were assayed with a high-sensitivity epinephrine ELISA kit (BAE-5100R, Labor Diagnostika Nord, Nordhorn, Germany). Cortisol levels were measured with a competitive ELISA kit (EIAHCOR, Invitrogen, Waltham, MA, USA) and corticosterone levels were measured with a corticosterone ELISA kit (RE52211, IBL, Hamburg, Germany).

### Flow cytometry

Isolated neutrophils (20 nM PMA-treated) or splenocytes were washed and resuspended in FACS buffer (1% BSA/PBS). Receptor expression was assessed with Ly6G (551461, BD Biosciences, San Jose, CA, USA) β2-AR (R11E1) Alexa Fluor 647 (L1824, Santa Cruz, Dallas, TX, USA) and GR (G-5) FITC (sc-393232, Santa Cruz, Dallas, TX, USA). Data were collected on a Cytek Aurora (30,000 events/sample) and analyzed with SpectroFlo after spectral unmixing.

### Open field test

Approximately 2.5 hours after dosing, the Open Field Test (OFT) was conducted to assess exploratory locomotor behaviors and depressive-like behaviors in a novel environment as previously described (45). Each mouse was placed at the center of a 421 mm x 421 mm x 300 mm (LWH) Plexiglas box under dim red-light illumination and recorded from above for 5 minutes. Between trials, the arena was wiped clean with 70% ethanol. Videos were processed using ANY-maze 7.44 software to measure total distance traveled around the open field, average speed of travel, time spent immobile, and number of center crossings. Immobile episodes were counted when no motion was observed for > 5 seconds. Center crossings were counted when motion was detected through the central zone (25% of the arena). Two-way ANOVA was conducted in GraphPad Prism (GraphPad Software, San Diego, CA, USA) 10.3.0 to compare the main effects and interaction of treatment and sex.

### Analysis of published datasets

Normalized transcriptome datasets from blood of healthy subjects and patients with MDD were extracted from the series GSE46743 and GSE98793 available on the Gene Expression Omnibus (GEO) database. The extracted data was used for gene expression analysis.

### Statistical analysis

Data were analyzed in GraphPad Prism 10.5.0. Results are presented as mean ± SEM. Normality was tested with Shapiro–Wilk. Two-group comparisons used unpaired *t* tests. Multi-group comparisons used two-way ANOVA (regular or RM) with Tukey’s post hoc test where appropriate. Statistical significance was set at *p* < 0.05 (*p* < 0.05, \**p* < 0.01, \*\**p* < 0.001).

## Results

### Chronic Restraint Stress induces NET formation

To investigate the effect of stress on NET formation, we subjected mice to chronic restraint stress (CRS) 3h every day for 28 days (Fig. 1A). The restraint stress model is a proven model of psychological stress characterized by activation of the HPA axis and SNS, and behavior changes including depressive-like behavior (46). On day 29, mice were sacrificed, and the plasma was collected and corticosterone levels and NETs levels in plasma were determined. The levels of corticosterone in plasma was similar between control and CRS mice on day 0 but showed a persistent increase up to day 21 (Fig. 1B). Intriguingly, CRS led to increased levels of circulating NETs as measured by extracellular DNA and neutrophil elastase in the plasma (Fig. 1C, D).

**Fig. 1.**
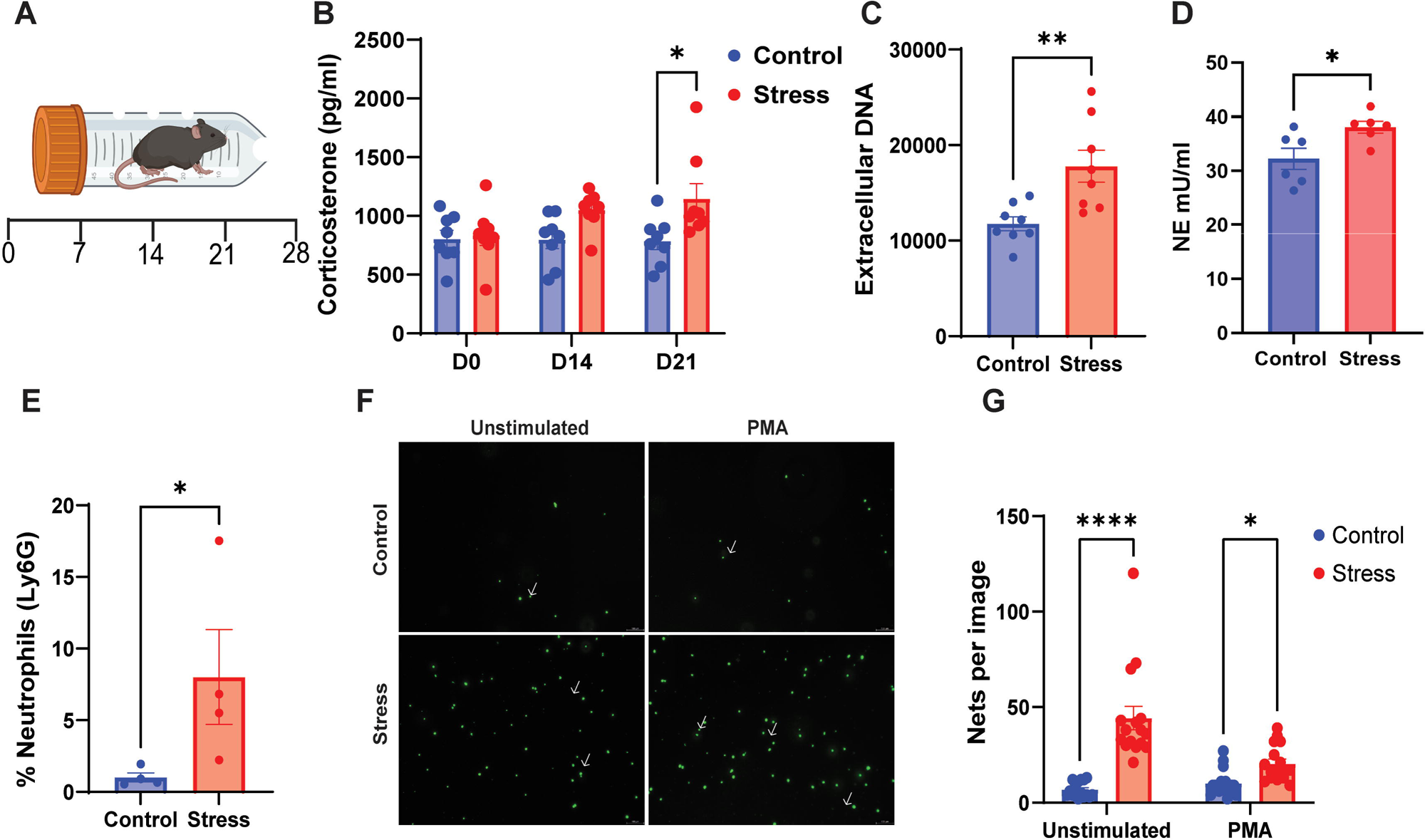
Chronic restraint stress promotes NET formation. (A) Schematic illustration of the chronic restraint stress (CRS) model in mice. (B) Plasma corticosterone levels measured at baseline (day 0), day 14, and day 21 in control and CRS mice. (C) Plasma NET levels quantified as extracellular DNA or elastase levels (D) at day 28. (E) Splenic neutrophil infiltration at day 28. (F-G) NET-forming capacity of blood neutrophils isolated from control and CRS mice under basal and PMA-stimulated conditions. White arrows depict NETs. Data represent mean ± SEM (n = 5 mice/group). *p < 0.05, **p < 0.01, ****p < 0.0001.

Previous studies have established stress-induced blood neutrophilia (28, 47). Therefore, we tested whether stress also led to neutrophil expansion in the lymphoid organs by flow cytometry. CRS led to increased neutrophil mobilization to the spleen (Fig. 1E). Given that chronic stress reprograms myeloid cells to a hyperresponsive inflammatory state (48) we explored the possibility that chronic stress will induce greater NET forming potential in neutrophils. We isolated neutrophils from the blood of CRS and control mice and activated them to form NETs. CRS led to increased NET formation compared to control (Fig. 1F-G). Thus, CRS leads to increased levels of NETs in the plasma and reprograms neutrophils to a hyperinflammatory state marked by a heightened ability to form NETs.

### Stress hormones directly induce NET formation

Given that physiological stress induces NET formation (Fig. 1C), we tested whether cortisol and epinephrine produced during stress will directly induce NETs. We isolated neutrophils from the blood of healthy volunteers and cultured them in the presence of varying concentrations of cortisol. Cortisol at 0.5mM did not lead to NET formation, however, higher levels of cortisol resulted in NETs formation (Fig. 2A-C). Interestingly, cortisol synergistically increased PMA-induced NETosis (Fig. 2C) supporting our previous observation that stress reprograms neutrophils to a hyper-inflammatory state marked by increased NET formation. Similarly, epinephrine induced NET formation and enhanced the NET-forming ability of neutrophils (Fig. 2D-F). Thus, both epinephrine and cortisol directly contribute to increased NET formation following physiological stress.

**Fig. 2.**
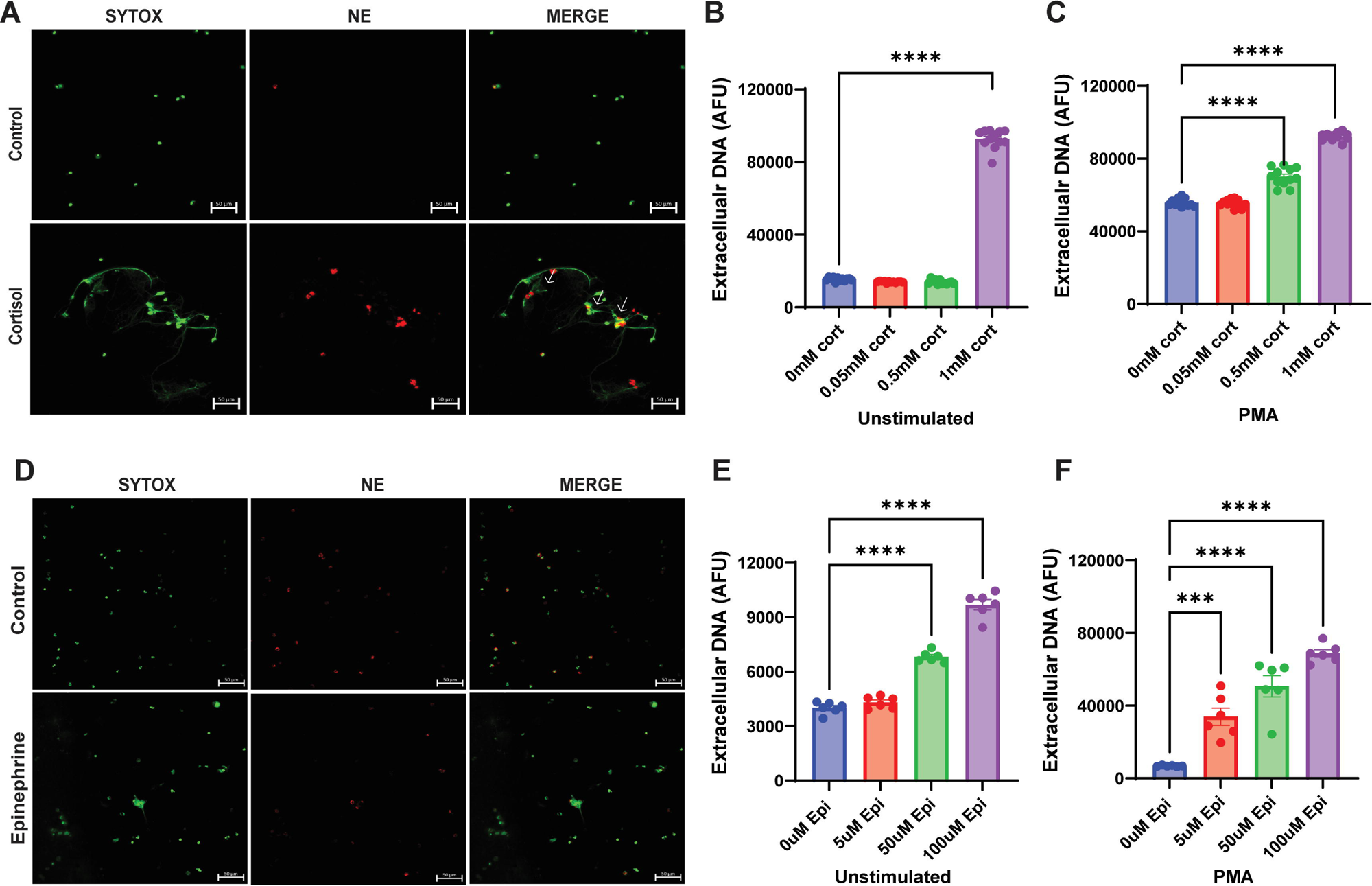
Stress hormones induce NET formation. Neutrophils purified from human peripheral blood were cultured with varying concentrations of cortisol (A-C) or epinephrine (D-F) with or without PMA activation. NETs released in culture were quantified by sytox green fluorescence. NETs were visualized by confocal microscopy as the colocalization of extracellular DNA and neutrophil elastase. White arrows depict NETs. ***p < 0.001, ****p < 0.0001.

### Bidirectional crosstalk between neutrophils and neuroendocrine-autonomic stress response

Previous studies have shown that activated neutrophils produce epinephrine which contributes to neutrophil inflammatory response (39). Increased production of stress hormones have been reported in patients with psychiatric diseases with a concomitant increase in the expression of adrenergic and glucocorticoid receptors on neutrophils (49, 50) suggesting that stress hormones can directly modulate neutrophil function. We therefore tested whether neutrophils produce the stress hormones epinephrine and cortisol during NETs formation. We activated neutrophils with PMA to induce NETs and analyzed the supernatant for epinephrine and cortisol production by ELISA. We found that NET-forming neutrophils also release the stress hormones epinephrine and cortisol (Fig. 3A, B). Interestingly, neutrophils activated to form NETs upregulate the expression of adrenergic and glucocorticoid receptors (Fig. 3C-F). Thus, our data indicates a bi-directional neuroimmune pathway whereby immune activation amplifies glucocorticoid and adrenergic signaling contributing to mood changes as seen in ‘sickness behavior’ (51) and highlights a feed forward mechanism whereby stress hormones induce NET formation and NET formation in turn, amplifies signaling by stress hormones thereby perpetuating neuro-immune pro-inflammatory interactions.

**Fig. 3.**
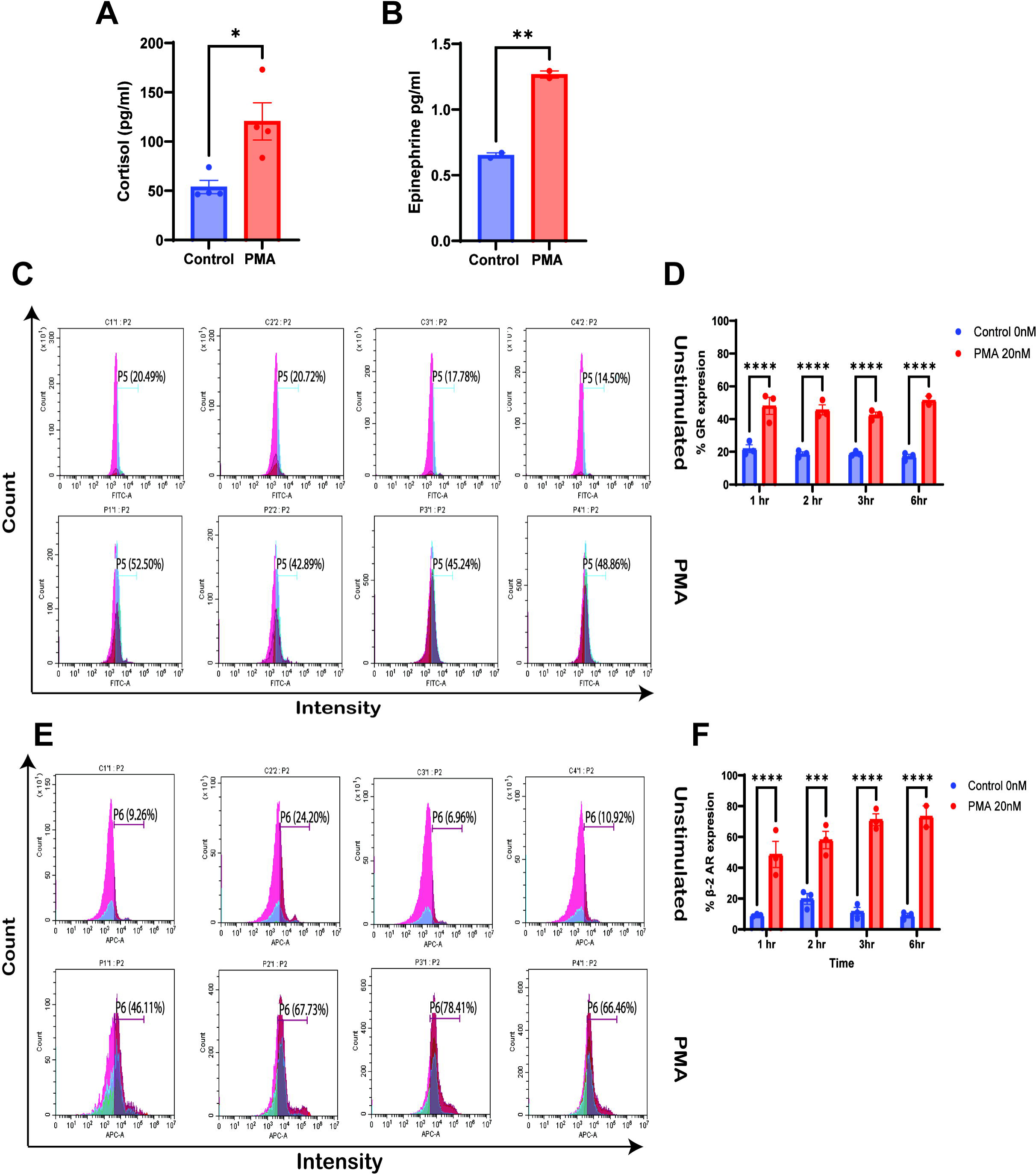
NET-forming neutrophils release stress hormones and upregulate glucocorticoid receptor (GR) and β2-adrenergic receptor (β2AR) expression. (A-B) ELISA quantification of cortisol levels (A) and epinephrine levels (B) in culture supernatants of neutrophils stimulated with PMA versus untreated controls. (C–D) Flow cytometry analysis of surface GR expression at 4 h post-PMA stimulation. (E–F) Flow cytometry analysis of β2AR expression at 4 h post-PMA stimulation. Data represent mean ± SEM. *p < 0.05, **p < 0.01, ****p < 0.0001.

### NETs contribute to behavioral changes in mice

NETs have been shown to be elevated in patients with schizophrenia (44) but whether NETs contribute to behavioral changes in these patients is unclear. Therefore, we tested the ability of NETs to contribute to behavioral changes in mice. We injected NETs into mice and investigated the ability of NETs to induce depressive-like behavior using the open field test (OFT). We used LPS as positive control, because LPS has been previously shown to induce depressive-like behavior in mice (52). Strikingly, similar to LPS, NETs administration led to decrease in exploratory behavior and increased depressive-like behavior in the animals in the OFT (Fig. 4 A-D). Our data indicates that NETs induced by stress could contribute to behavioral disorders.

**Fig. 4.**
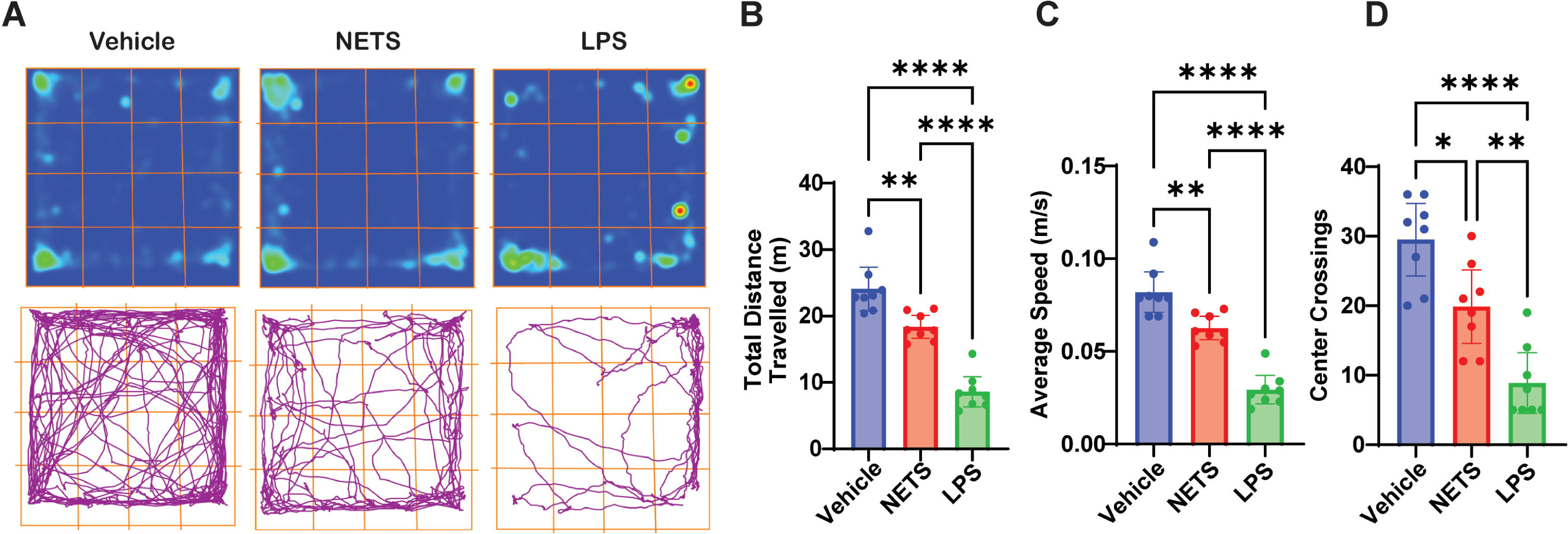
NET injection induces depressive-like behavior in mice. (A) Representative heat maps of locomotor activity in vehicle-, NET-, and LPS-exposed mice. (B) Total distance travelled in the open field. Average speed (C) and number of center crossings (D) in the open field. Data represent mean ± SEM (n = 5 mice/group). *p < 0.05, **p < 0.01, ****p < 0.0001.

### Activation of glucocorticoid receptor upregulates NET genes in humans

NET formation proceeds through the action of the enzyme peptidylarginine deiminase 4 (PAD4) which promotes the citrullination of histones leading to chromatin decondensation, rupture of the nuclear envelope and the release of NETs (53). During NET formation, neutrophils release their granular contents including neutrophil elastase and myeloperoxidase (MPO) into the extracellular space (53). Our data shows that stress induces NET formation in mice (Fig. 1C, D). We therefore investigated whether activation of the glucocorticoid receptor in humans leads to NET formation by analyzing gene expression changes of *ELANE*, *MPO* and *PADI4* in patients with major depressive disorder (MDD) and control group at baseline and following an oral administration of the glucocorticoid agonist dexamethasone using publicly available datasets from the GEO database (GSE46743). At baseline, MDD patients had a higher trend of gene expression for NET proteins though this was not significant (Fig. 5A-C). However, both control group and MDD group showed elevated expression of *ELANE*, *MPO* and *PADI4* following oral administration of dexamethasone (Fig. 5A-C). We further analyzed another dataset of a cohort of MDD patients (GSE98793) and found that at baseline, there was significantly elevated levels of *ELANE*, *MPO* and *PADI4* in the blood of patients with MDD (Fig. 6). Taken together, our findings highlight the role of stress in driving NET formation which contributes to psychopathology in patients with MDD.

**Fig. 5.**
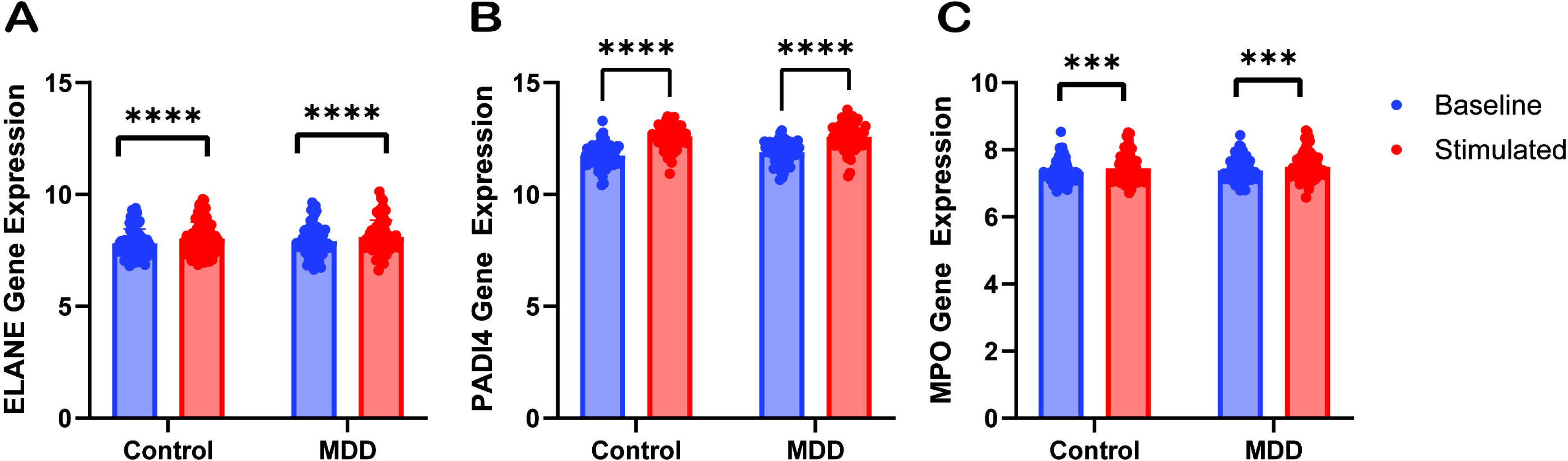
Activation of glucocorticoid receptor increases gene expression of NET proteins in humans. (A-C) Relative expression of blood *ELANE*, *MPO* and *PADI4* in MDD patients and healthy controls at baseline and following oral administration of dexamethasone. Data from GEO GSE46743. ***p < 0.001.

**Fig. 6.**
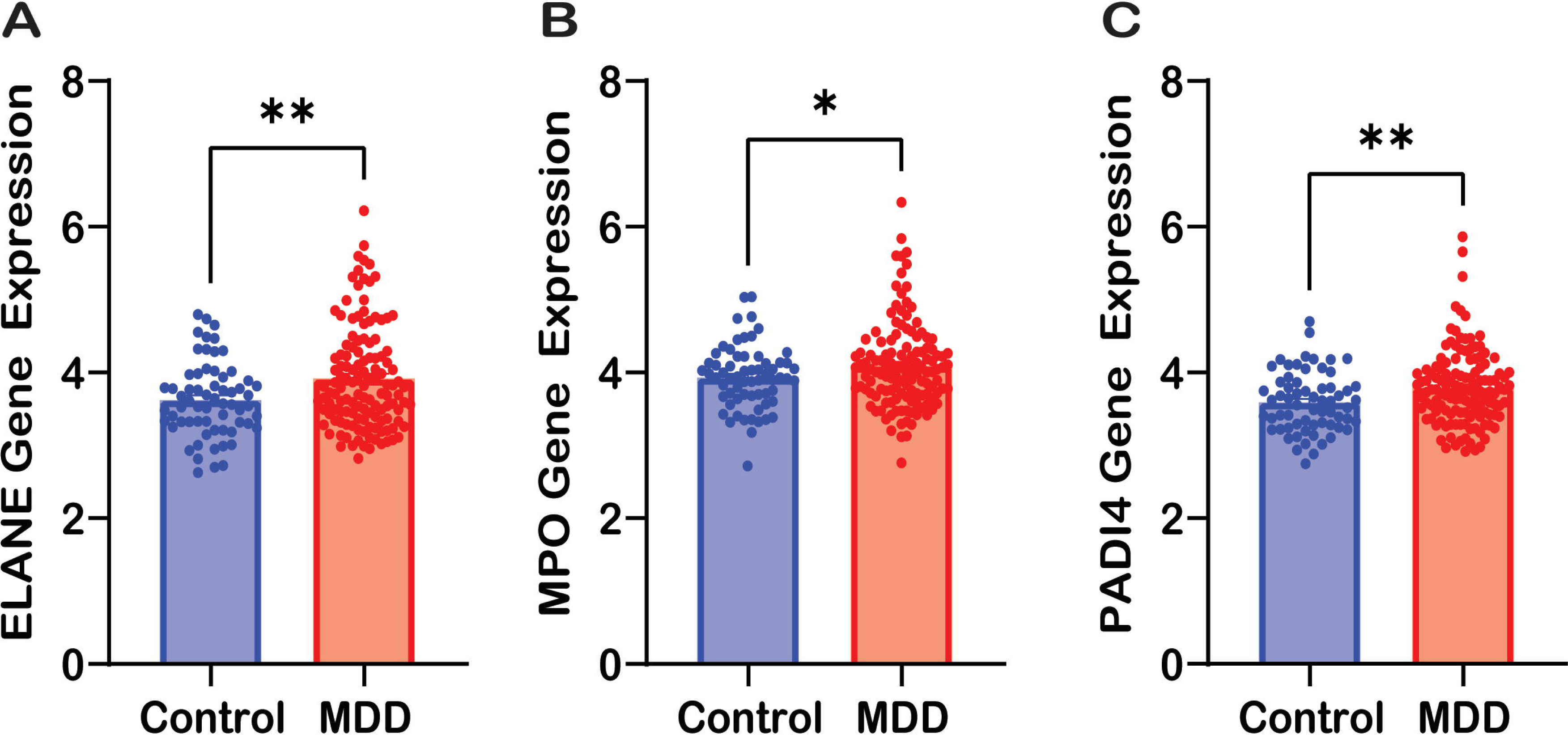
Elevated gene expression of NET proteins in blood is associated with depression in humans. A-C. Relative expression of blood *ELANE*, *MPO* and *PADI4* in MDD patients and healthy individuals. Data from GEO GSE98793. *p < 0.05, **p < 0.01.

## Discussion

There is compelling evidence that neuro-immune interactions contribute to psychopathology but the underlying mechanisms remain unclear. It is therefore pertinent to elucidate the effect of stress hormones on immune function. Glucocorticoids are generally recognized as immune-suppressive agents and are available in the clinic as anti-inflammatory drugs (54). However, given that physiological stress has been shown to promote systemic inflammation (22, 55), it is conceivable that stress hormones contribute to the inflammatory response. Indeed, both cortisol and epinephrine have been shown to promote the inflammatory response of immune cells (56, 57).

Neutrophils are the most abundant leukocytes in the body and are the first cells to arrive at the site of acute inflammation (58). Physiological stress has been shown to lead to neutrophil expansion and activation (47). Indeed, among all leukocytes, neutrophils have the most sustained increase in numbers following physiological stress and continue an upward trajectory even when the numbers of other immune cells were decreasing (28). Hence, there is significant interest in the effect of stress hormones on neutrophil functions. Importantly, neutrophils have been implicated in the development of mood disorders. Indeed, neutrophil-lymphocyte ratio increases in patients with MDD (59, 60) and neutrophils are the major immune subset most predictive of symptom severity in MDD (61).

Since, their discovery as a unique mechanism of neutrophil pathogen killing, NET formation has been shown to be a double-edged sword (62) and inflammation driven by NET formation has been shown to contribute to the pathogenesis of several diseases including rheumatoid arthritis (63), sepsis (41), lupus (64), cancer (65) and COVID-19 (42). Since stress has been shown to lead to neutrophil expansion (28), it is conceivable that stress hormones induce NET formation and that stress-induced NETs will contribute to the pathogenesis of mental health disorders.

Here, we show that chronic restraint stress resulted in increased circulating NETs in mice and reprograms neutrophils to a hyper-NET forming state (Fig. 1). Also, stress hormones induce NET formation and activated neutrophils produce the stress hormones cortisol and epinephrine thereby highlighting an autocrine/paracrine mechanism for the modulation of neutrophil inflammatory responses by stress hormones (Fig. 2). This is further supported by our finding that induction of NETs formation in neutrophils leads to increased expression of glucocorticoid and adrenergic receptors indicating that neutrophil activation can perpetuate local and systemic activation of the HPA axis and the sympathetic nervous system (Fig. 3). Furthermore, we show that genes encoding NET proteins are increased in stressed individuals (Fig. 5) and in patients with major depressive disorder (Fig. 6). Our data adds to the growing body of evidence that neutrophils and NETs are implicated in psychopathology.

How do stress hormones contribute to NET formation? NET formation is dependent on the activation of neutrophil pro-inflammatory pathways and NADPH oxidase (NOX) activity (53). The activation of NOX leads to the production of reactive oxygen species (ROS) (53). ROS facilitates the release of proteases such as neutrophil elastase from cytoplasmic granules which degrade histones and promotes NETs formation. ROS production further activates signaling cascades that ultimately promote histone citrullination by PAD4 and rupture of the nuclear envelope leading to NET release (33). It is possible that stress hormones induce ROS production by neutrophils and thus leading to NETs formation. Indeed, a recent study showed that glucocorticoids induce NETs formation in a ROS-dependent manner (66).

A major finding of our study is that injection of NETs leads to depressive-like behavior in mice (Fig. 4). To our knowledge, this is the first observation of NET-induced behavioral changes in an experimental setting. It remains to be determined how NETs contribute to changes in behavior. Pro-inflammatory molecules like LPS are known to induce behavioral changes in mice (52). Hence, NET components including elastase and histones which are proinflammatory can also induce alterations in behavior in mice. The mechanisms by which NETs contribute to behavior changes remains the subject of investigation in our laboratory.

Our work suggests that NETs production by neutrophils could contribute to the pathogenesis of psychiatric diseases. Indeed, Corsi-Zuelli et al showed that increased plasma NETs have been reported in patients with schizophrenia and this increase was shown to be associated with early life adversity (44). Kong et al, also showed that NETs contribute to depressive state in mice (52). Another study recently showed that catecholamines released during chronic stress promote NET formation and exacerbates cerebral amyloid angiopathy (67). In summary, our work shows that stress hormones modulate neutrophil function including NET formation and that stress-induced NETs could contribute to the development of psychopathology.

## Acknowledgement

E.O was supported by funding from Northeastern University.

K.T was supported by Postdoctoral Fellowship from the College of Science, Northeastern University.

